# ARGformer: learning on ancestral recombination graphs with transformers

**DOI:** 10.64898/2026.02.11.705405

**Authors:** David Bonet, Cole Shanks, Marçal Comajoan Cara, Jordi Abante, Alexander G. Ioannidis

## Abstract

Recent advances in inference of the ancestral recombination graph (ARG), which describes how segments of chromosomes trace back through recombination and shared lineages, have made it possible to reconstruct genome-wide genealogies for large cohorts, but it remains difficult to summarize and use this information for population genetic analyses. We present ARGformer, an encoder-only transformer that learns context-dependent embeddings with a self-supervised masked objective finetuned with contrastive learning for downstream retrieval tasks. We train ARGformer on genealogies from coalescent simulations and on genealogies inferred from ancient and present-day *Homo sapiens* genomes. Using only these learned embeddings, without access to genotype matrices, ARGformer captures patterns of global population structure and supports ancestry inference through clustering and nearest-neighbor retrieval. On genealogies that include archaic hominins, ARGformer can highlight Denisovan-derived segments in Oceanian genomes and reveals Oceanian-like ancestry in South American Indigenous populations. ARGformer is available at https://github.com/AI-sandbox/ARGformer.

## 1. Introduction

The high dimensionality of genomic data, especially in large population-scale sequencing studies such as national biobanks, has motivated the development of a variety of methods for visualizing population structure (Bycroft et al., 2018). The most popular of these is principal component analysis (PCA) (Patterson et al., 2006), as well as uniform manifold approximation and projection (UMAP) (Diaz-Papkovich et al., 2019). In the context of population genetics, these dimensionality reduction techniques provide insight into population structure and relatedness. Deep learning has been applied for visualizing population structure in the form of variational autoencoders (VAEs) trained on SNP genotypes. VAEs produce low-dimensional latent embeddings that capture nonlinear population structure and often preserve global geometry better than t-SNE or UMAP applied directly to SNP genotypes (Battey et al., 2021; Geleta et al., 2026). Other approaches include MAPS, which estimates time-stratified migration-rate surfaces from identity-by-descent (IBD) tracts, providing temporal visualization of structure (Al-Asadi et al., 2019). Collectively, these methods provide powerful low-dimensional summaries of genetic data, but they operate directly on genotypes or derived summaries, rather than on the underlying genealogies that generate those genotypes.

Patterns of allele sharing and population structure are embedded in the history of genealogies under the coalescent with recombination (Hudson, 1983; Griffiths & Marjoram, 1997). The ancestral recombination graph (ARG) provides a unifying representation of this history by encoding the collection of local genealogical trees along the genome together with recombination events that rearrange ancestry through time (Griffiths & Marjoram, 1997; Lewanski et al., 2024; Nielsen et al., 2025). Genealogical interpretations of principal component analysis make this link explicit by viewing genotype structure as a projection of the underlying genealogy (McVean, 2009). In the presence of recombination, the ARG encodes information about population size changes, migration and admixture, recombination and mutation processes, linkage disequilibrium, and identity by descent (Nielsen et al., 2025). In principle, methods that learn directly from ARGs have the potential to unify many classical population genetic analyses within a common representation.

Recent methods can now infer genome-wide genealogies at scale for large cohorts, demonstrating the utility of inferred ARGs for genome-wide association studies and population structure analyses (Nielsen et al., 2025; Rasmussen et al., 2014; Speidel et al., 2019; Kelleher et al., 2019; Zhang et al., 2023; Gunnarsson et al., 2024; Deng et al., 2025). However, there is not yet a standard, scalable self-supervised framework for representation learning directly on genome-wide genealogies or ARG-derived structures analogous to the way that VAEs have been applied to genotypes. This could be related to the challenges associated with such a task. For example, it is not immediately clear how to input an ARG into a deep learning model in the way that VAEs can process genotype data. In addition, the complete ARG can be an ultra-large graph even for a modest cohort, and the marginal coalescent trees also become very large graphs for medium to large scale cohorts. More generally, although deep learning has been used for population genetic inference and for learning features from sequence data (Sheehan & Song, 2016; Flagel et al., 2019; Schrider & Kern, 2018), there is no standard framework for self-supervised representation learning directly on genome-wide genealogies. Recently, transformer architectures have begun to be applied to population genetics. A recent example is cxt, a decoder-only transformer that autoregressively predicts pairwise coalescence times from local mutational context (Korfmann et al., 2025). This line of work shows that transformers can be effective for genealogically informed inference from sequence-derived inputs. ARGformer addresses a complementary problem: rather than inferring coalescence from mutational context, it learns representations directly from inferred genealogies, enabling downstream tasks such as visualization, clustering, and retrieval on ARG-derived structures.

An architecture for efficiently encoding sequences in natural language processing has been encoder-only transformers, and in particular Bidirectional Encoder Representations from transformers (BERT) models (Devlin et al., 2019). Masked language modeling objectives allow such models to learn rich, context-dependent representations in a fully self-supervised fashion, and recent work has introduced more efficient encoder-only transformers that scale to large training corpora (Warner et al., 2025). Encoder-only transformers and masked prediction objectives have also been successfully transferred to a wide range of biological sequence and graph domains, where they provide general-purpose embeddings for downstream tasks (Rives et al., 2021; Ji et al., 2021). More broadly, recent theoretical and empirical work has shown that pure transformer architectures, with suitable tokenization schemes, can act as powerful and competitive graph learners, often rivaling or surpassing message passing graph neural networks (Kim et al., 2022).

Here we explore an encoder-only model for representing coalescent events in the ARG, which we call ARGformer. The model encodes, for each extant haplotype, the path from the leaf to the root as a token sequence, exploiting shared coalescent events across paths. ARGformer learns context-dependent embeddings that recapitulate global population structure, capture local genealogical relationships, and support downstream local ancestry analyses via clustering or nearest-neighbor retrieval in embedding space. The advantage of ARGformer is that it compresses the massive, ultra-high-dimensional structure of the ARG into a dense, accessible latent space, enabling downstream methods to query complex coalescent histories without directly parsing the raw graph topology. Conceptually, our approach is related to genealogical interpretations of principal component analysis in population genetics, which view genotype structure as a projection of the underlying genealogy (McVean, 2009), but ARGformer operates directly on the topological structure of inferred genealogies, leveraging the evolutionary history already captured by the ARG rather than attempting to learn these relationships from raw genotype matrices.

In applications to both simulated genealogies and ARGs inferred from ancient and present-day *Homo sapiens* genomes, we show that ARGformer embeddings alone, without access to genotype matrices, recover patterns of global population structure and support ancestry inference via clustering and nearest-neighbor retrieval. On ARGs that include archaic hominin genomes, ARGformer embeddings highlight Denisovan-like segments in Oceanian genomes, consistent with previous observations of Denisovan introgression in Papuans (Jacobs et al., 2019; Vespasiani et al., 2022), and reveal Oceanian-like ancestry patterns present in some Indigenous South American populations that are not evident in the other Indigenous American groups. This echoes previously observed Australasian-related ancestry in Amazonian groups (Skoglund et al., 2015; Castro e Silva et al., 2021). These examples illustrate how local genealogies along the genome encode complex demographic and admixture histories, and how representation learning on ARGs can make this information accessible for visualization and downstream analyses.

## 2. Materials and methods

### 2.1. BERT model for encoding paths in ancestral recombination graphs

There are many possibilities for encoding ARGs with a transformer. Rather than encoding an entire ARG or full marginal coalescent trees, we encode the coalescent events along leaf-to-root paths. Because different paths share internal ancestral nodes and adjacent marginal trees share substantial structure along the genome, this representation retains substantial genealogical context while remaining scalable. This approach is similar to the sub-sampling also seen in representation learning for graphs and enables better scalability (Hou et al., 2022). ARGformer encodes paths of marginal coalescent trees and augments them with positional encodings that reflect the ordering of coalescence events and the local tree topology along the path, providing a flexible representation model for ARGs. Each node in the inferred genealogy has a unique identifier. Because marginal trees share ancestral nodes, and adjacent trees along the genome share topologies between recombination events, encoding these paths preserves coalescent and recombination history. By learning from repeated co-occurrence of ancestral nodes across samples and genomic intervals, the model can capture genealogical relationships and their evolution through time along the genome.

ARGformer is an encoder-only transformer that operates on these path sequences to learn vector representations of ARG nodes and haplotypes (Figure 1). During self-supervised pretraining, we adopt a masked-node objective: we randomly mask node tokens along each path and optimize a cross-entropy loss to predict the masked nodes, analogous to masked language modeling but applied to the ARG. Concretely, let a leaf-to-root path be a sequence of tokens **x** = (*x*_1_, …, *x*_L_), where each token is a node identifier. Let *M* ⊆ {1, …, *L*} denote the set of masked positions, sampled by independently including each position *i* in *M* with probability *p*_mask_ = 0.30 (Warner et al., 2025). Let 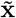 be the corrupted input sequence obtained by replacing *x*_i_ with a special `[MASK]` token for all *i*∈ *M* . Given the transformer hidden state 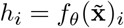 at position *i*, we predict the original token via a softmax head over the vocabulary

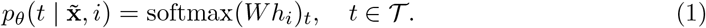

**Figure 1:**
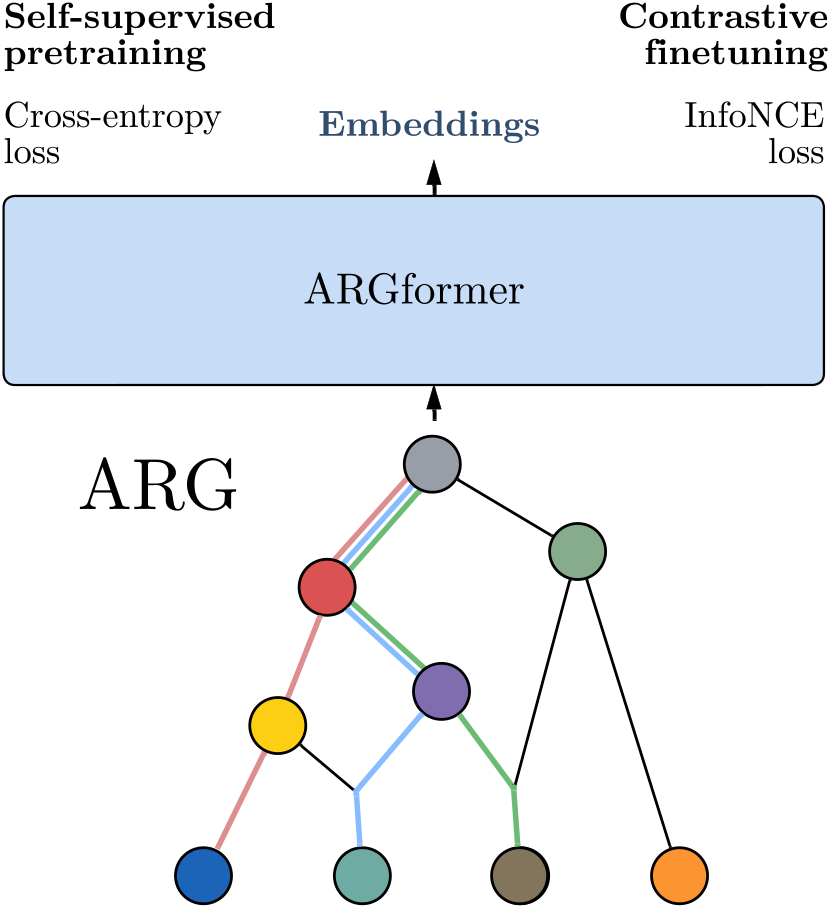
Overview of ARGformer. Schematic ARG, with colored edges showing example marginal paths, which are passed through ARGformer, an encoder-only transformer, with self-supervised masked pretraining objective, and contrastive finetuning for downstream usage of learned embeddings (e.g., visualization, clustering, and retrieval).

We then minimize the masked cross-entropy loss averaged over masked positions,

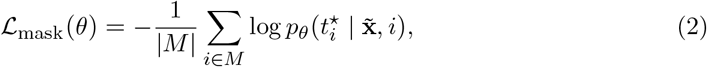

where 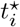 is the ground-truth token at position *i*. We build ARGformer on top of a

ModernBERT-style encoder that incorporates recent architectural and optimization improvements, which substantially improves training efficiency on large collections of data (Warner et al., 2025).

Although the resulting model after pretraining is already infused with global population structure, we finetune the pretrained encoder for downstream tasks with a supervised contrastive objective that pulls together embeddings of sequences sharing the same reference population label while pushing apart embeddings from different labels within each minibatch.Given a batch of *B* labeled sequences 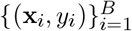 with labels *y*_i_ ∈ {1, …, *C*}, we encode each sequence with the transformer backbone followed by a pooling head to obtain embeddings:

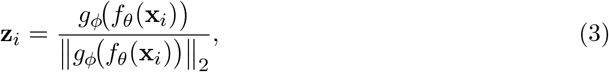

where *f*_θ_ is the transformer encoder, *g*_*ϕ*_ is the pooling head, and embeddings are *l*_2_-normalized. For each anchor *i*, define the in-batch positive set *P* (*i*) = {*j* ≠ *i* : *y*_j_ = *y*_i_} . We minimize the supervised contrastive (InfoNCE-style) loss (Oord et al., 2018)

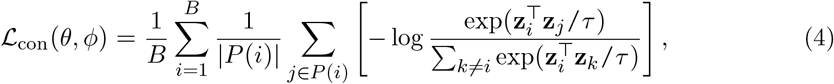

where *τ >* 0 is a temperature hyperparameter (default *τ* = 0.05). The denominator sums over all in-batch samples except the anchor, providing implicit hard negatives from other classes. Anchors with no same-class partners in the batch ( *P*| (*i*) | = 0) are excluded from the loss. To mitigate class imbalance, we apply inverse-frequency weighting:

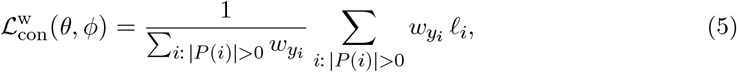

where *l*_i_ denotes the per-anchor term in ℒ_con_, and 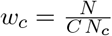 with *N* the total number of train examples and *N*_c_ in class *c*.

### 2.2. ARGformer on simulated data

We first applied ARGformer to simulated data generated using `msprime` (Kelleher et al., 2016). We simulated 1600 samples from four populations (Admixed=1000, European=200, African=200, East Asian=200) in the three-population out-of-Africa demography (AmericanAdmixture 4B18) from the `stdpopsim` catalog (Adrion et al., 2020), for a single 30 million base pair (bp) chromosome, using a flat recombination rate of 1×10−8 per bp generation and a mutation rate of 1.25e-8 per bp per generation (Lauterbur et al., 2023; Browning et al., 2018). We generated a dataset for pretraining using `tskit` to extract coalescent events (Wong et al., 2024). To better evaluate ARGformer on inferred ARGs, we inferred the ARG using `tsinfer+tsdate` on the simulated genotypes, without phasing or genotype error (Kelleher et al., 2019). We split this dataset into training (90%) and validation (10%) for pretraining.

### 2.3. Application of ARGformer to an inferred ARG of ancient and modern DNA

To evaluate the model on empirical data, we utilized the inferred genealogy of ancient and present-day human genomes from (Wohns et al., 2022). From this ARG, we constructed two task-specific datasets designed to probe complex evolutionary and admixture histories. Unique marginal paths for these datasets were split 90/10 into train and validation sets, and full population composition and dataset statistics are provided in Supplementary Section S1.

## 3. Results

We first evaluated ARGformer on coalescent simulations to determine whether embeddings recover patterns of global population structure and support downstream local ancestry inference. Crucially, we also examined the intrinsic properties of the learned latent space, demonstrating that the self-supervised embeddings inherently capture genealogical depth. Specifically, predicting the number of coalescent events along a path, and that the model’s attention heads specialize to distinct genealogical motifs. Building on these foundational capabilities, we then applied ARGformer to the ARGs inferred from ancient and present-day *Homo sapiens* genomes. Relying solely on the learned embeddings, we localized Denisovan ancestry in Oceania and identified Oceanian-like affinities within specific Indigenous South American populations. Finally, we summarize model behavior, convergence, and computational efficiency during self-supervised pretraining (Supplementary Section S2).

### 3.1. Capturing population structure

ARGformer embeddings provide a chromosomal segment-level view of population structure derived from local genealogies (Figure 2). In the genotype PCA (Figure 2A), each point aggregates information across the entire genome, so individuals with contributions from multiple ancestries will fall between the three continental clusters. In contrast, each point in Figure 2B and C corresponds to a single marginal-path embedding, representing a local genealogy segment in the genome. Under the simulated admixture demography, local segments predominantly trace to one of the unadmixed sources and therefore concentrate within reference ancestry clusters.

**Figure 2:**
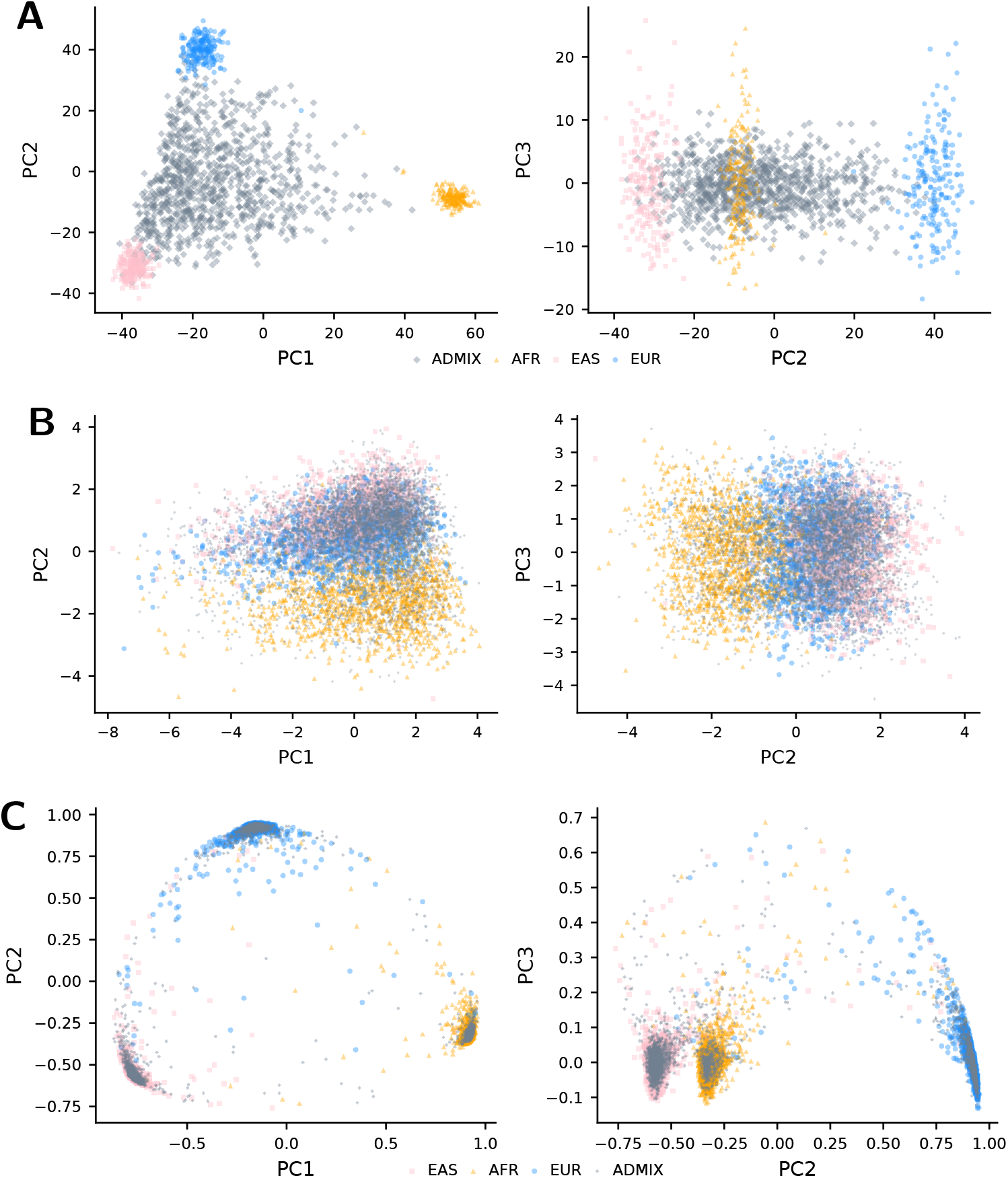
Population structure and local-ancestry geometry on simulated data. Colors indicate population label: East Asian (pink), African (orange), European (blue), and admixed (gray). **A**: PCA of ground-truth genotypes (points are individuals; PC1 vs PC2; PC2 vs PC3). PCA is fitted on reference populations (AFR, EAS, EUR), and ADMIX samples are projected. **B** and **C**: The same PCA protocol applied to ARGformer embeddings, which are generated without genotype matrices (points are haplotype marginal-path embeddings, multiple per individual). **B** shows the embeddings after self-supervised pretraining, while **C** shows the contrastive embeddings.

To evaluate the contribution of our distinct training stages, we performed an ablation comparing the model’s representations immediately after self-supervised pretraining (Figure 2B) to those obtained after contrastive finetuning (Figure 2C). This comparison shows that pretraining alone already captures inherent aspects of global population structure directly from the ARG topology, prior to the introduction of any ancestry labels. Self-supervised pretraining is important because it learns a general genealogical representation from shared ancestral-node structure before contrastive training embeds held-out query lineages relative to labeled references. The subsequent contrastive finetuning then builds upon this foundation to more sharply separate the distinct local-ancestry clusters.

To test whether these self-supervised embeddings encode genealogical structure rather than only downstream separability, we trained a ridge probe on frozen embeddings to predict the number of coalescent events along each path. In the American simulation, the probe achieves strong held-out agreement between predicted and observed path depth (Figure 3A). The same analysis also generalizes to inferred genealogies from ancient and present-day humans in the archaic-modern setting (Figure 3B), showing that the learned representation captures aspects of genealogical depth in both simulated and real ARGs. In each case, performance collapses under a permutation control in which the training labels are randomly shuffled before fitting the same ridge probe, preserving the embedding distribution and label marginal while destroying the embedding-label correspondence. This indicates that the signal is genuinely present in the learned embedding geometry. In the archaic-modern ARG, the parity plot also shows a concentration of short paths with few coalescent events, consistent with the long-branch structure of archaic lineages. Additional probe results for the Oceania-America analysis, along with attention interpretability analyses of the self-supervised model, are provided in Supplementary Section S4.

**Figure 3:**
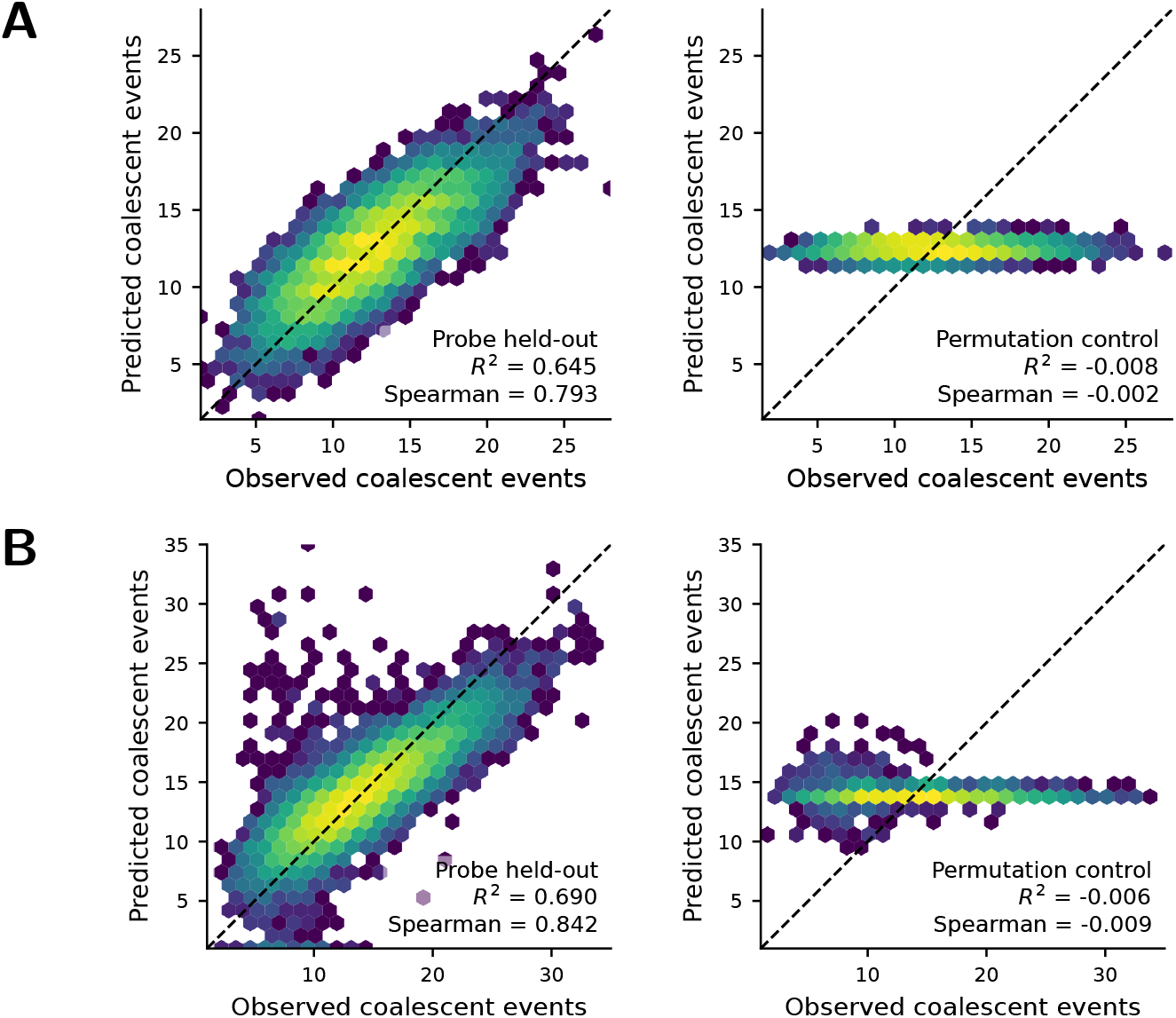
Self-supervised embeddings encode genealogical structure in both simulated and inferred ARGs. **A**: American simulation. **B**: Real archaic-modern inferred ARG. In each panel, the left plot shows held-out prediction of the number of coalescent events using a ridge probe on frozen embeddings, and the right plot shows the corresponding permutation control obtained by randomly shuffling the training labels before fitting the same probe.

Using only the learned ARG-based embeddings, we can separate local chromosomal segments without directly accessing genotype matrices. In this sense, ARGformer plays for genealogies and haplotypes a role that is analogous to PCA for genotypes, providing a low-dimensional visualization of population structure that is rooted in the inferred ARG.

We benchmarked local ancestry inference (LAI) in admixed individuals against FLARE (Browning et al., 2023). Because FLARE is a specialized local-ancestry method, we use it here as a task-specific baseline, whereas ARGformer is intended as a more general representation model for multiple downstream analyses on inferred genealogies. Using 10, 000 unique marginal paths sampled from admixed individuals, we considered two inference strategies based solely on ARGformer embeddings. The first performs PCA on embeddings of unadmixed reference individuals and retains the top 10 principal components. We then project each marginal path embedding into this 10-dimensional PC space and assign an ancestry label based on its cluster membership in this space (PCA clustering).

The second strategy treats each admixed marginal path as a query and retrieves its nearest neighbors among embeddings of reference haplotypes in embedding space, assigning ancestry labels based on the labels of the retrieved neighbors. In both cases we compare the resulting three-way local ancestry calls with those from FLARE, providing a quantitative benchmark for how well the embeddings capture population structure at the level of individual genomic segments. ARGformer achieved comparable accuracy to FLARE, a state-of-the-art local ancestry inference method and obtained better performance with the nearest-neighbor retrieval (Table 1).

**Table 1:** LAI precision and recall on simulated American admixture. FLARE is run on simulated genotypes, and ARGformer is run on the inferred ARG.

| Method | EUR |  | EAS |  | AFR |  |
| --- | --- | --- | --- | --- | --- | --- |
|  | Prec. | Rec. | Prec. | Rec. | Prec. | Rec. |
| FLARE | <u>98.41</u> | **98.54** | <u>98.85</u> | **99.74** | **99.70** | 96.70 |
| ARGformer PCA | **98.47** | 97.91 | 98.81 | <u>99.07</u> | <u>97.46</u> | <u>97.81</u> |
| ARGformer retr. | **98.47** | <u>98.00</u> | **98.88** | 99.04 | <u>97.46</u> | **97.90** |

### 3.2. Identifying Denisovan ancestry in Oceania

To test whether ARGformer embeddings capture archaic introgression signals without using genotype matrices, we framed archaic ancestry detection as a nearest-neighbor retrieval problem in embedding space. For each held-out population, we retrieved the top-5 closest haplotype-path embeddings and summarized the retrieved reference labels as a row-normalized distribution over reference populations, reporting both the raw vote share and a chance-corrected enrichment (Figure 4).

**Figure 4:**
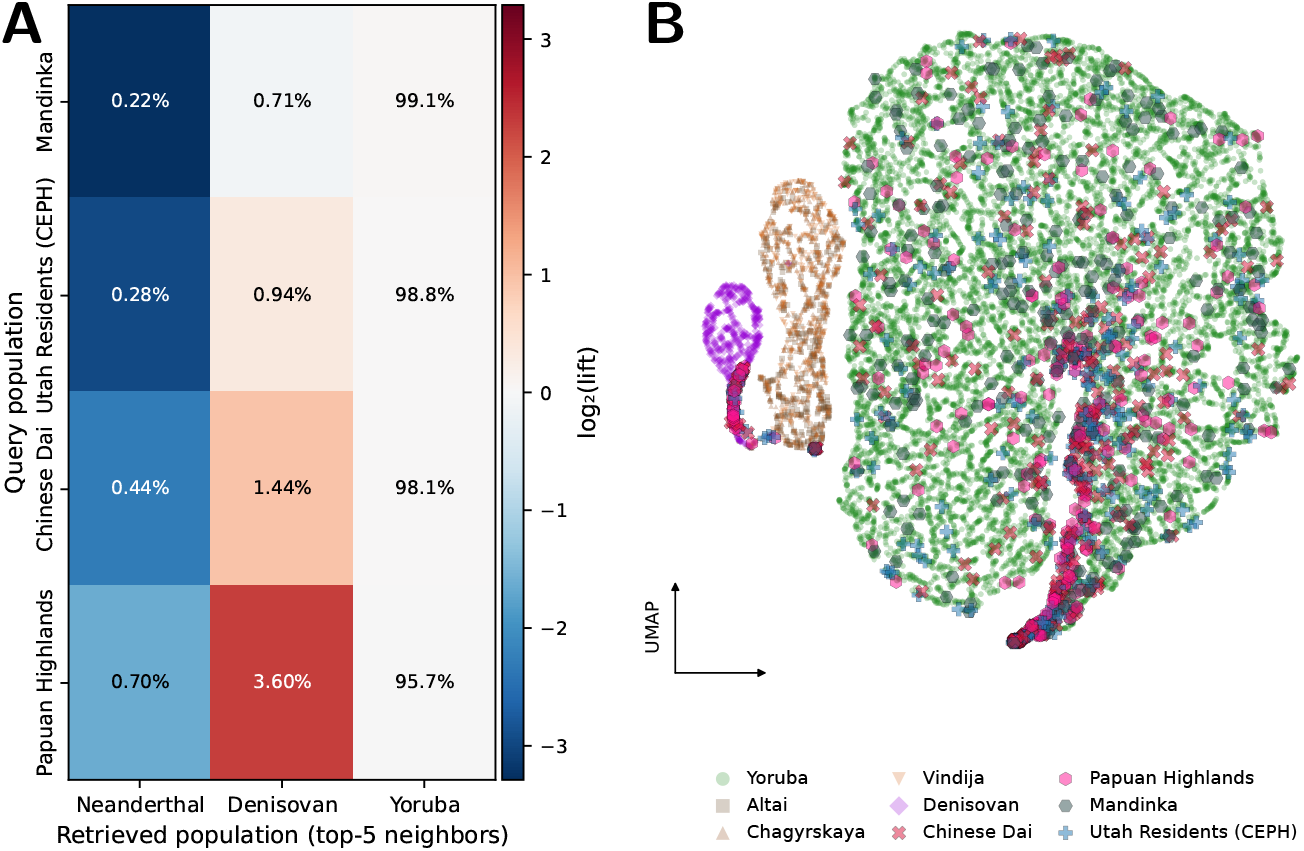
Denisovan segment retrieval in Oceania. **A**: Nearest-neighbor retrieval reveals Denisovan-enriched embedding neighborhoods in Oceanian haplotypes. For each held-out query haplotype (row), we retrieve its top-5 nearest neighbors and record the neighbors’ reference individual labels (columns). Each cell reports the row-normalized fraction of retrieved neighbors assigned to the corresponding reference population. Colors show the log-lift of retrieval relative to chance (log_2_(lift) = log_2_ 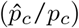where 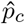 is retrieved vote share and *p*_c_ is corpus label frequency), so values above zero indicate enrichment beyond what would be expected from corpus composition. **B**: UMAP projection of haplotype-path embeddings for archaic references, modern population reference, and query populations.

Papuan Highlands stands out with a Denisovan neighbor fraction of 3.60% among its top-5 neighbors, greater than that of the comparison populations with non-Oceanian populations 1.44% (Chinese Dai), 0.94% (Utah Residents (CEPH) with Northern and Western European ancestry), and 0.71% (Mandinka) in the same analysis. This enrichment is reflected by the positive log-lift coloring in the Denisovan column (Figure 4A), where Mandinka is the only query population with a negative lift. In contrast, all query populations predominantly retrieve Yoruba-labeled neighbors (between 95.7% and 99.1% across rows), consistent with most genomic segments being closest to modern human variation. We also note that some non-Oceanian samples (e.g., Utah Residents (CEPH) with Northern and Western European ancestry) can retrieve a slightly higher fraction of Denisovan-labeled neighbors than Neanderthal-labeled neighbors. Because the archaic genomes were computationally phased without reference panels or imputation, and because contrastive finetuning can be sensitive to such technical artifacts, we treat these retrieval shares as relative enrichment signals rather than admixture proportions.

Together, this shows that ARGformer embeddings learned from inferred genealogies can localize Denisovan-like segments in Oceanian genomes, consistent with established Denisovan introgression in Papuan-related Oceanian populations (Jacobs et al., 2019; Vespasiani et al., 2022). Panel B in Figure 4 shows a UMAP projection, highlighting distinct clusters and a subset of query haplotypes falling on archaic clusters.

### 3.3. Oceania-like ancestry in South America

We next asked whether ARGformer embeddings capture the subtle Oceania-related affinities that have been previously reported in some Indigenous South American Amazonian populations using allele frequency based analyses (Skoglund et al., 2015; Castro e Silva et al., 2021). As in the archaic introgression analysis, we framed this question as a retrieval problem in embedding space: given a query set of marginal-path embeddings from Indigenous American populations, we retrieve the top-5 nearest neighbors of its haplotype-path embeddings from a reference corpus stratified into three regional groups (Americas, East Asian, and Oceania) and summarize neighbor identities as a row-normalized vote share, alongside a chance-corrected enrichment (Figure 5A). In this analysis, the Americas reference group comprises Pima, Maya, and Mixe individuals; the Oceania reference group comprises Australian, Bougainville, Papuan, Papuan Highlands, and Papuan Sepik; and the East Asian reference group comprises Han Chinese and Japanese.

**Figure 5:**
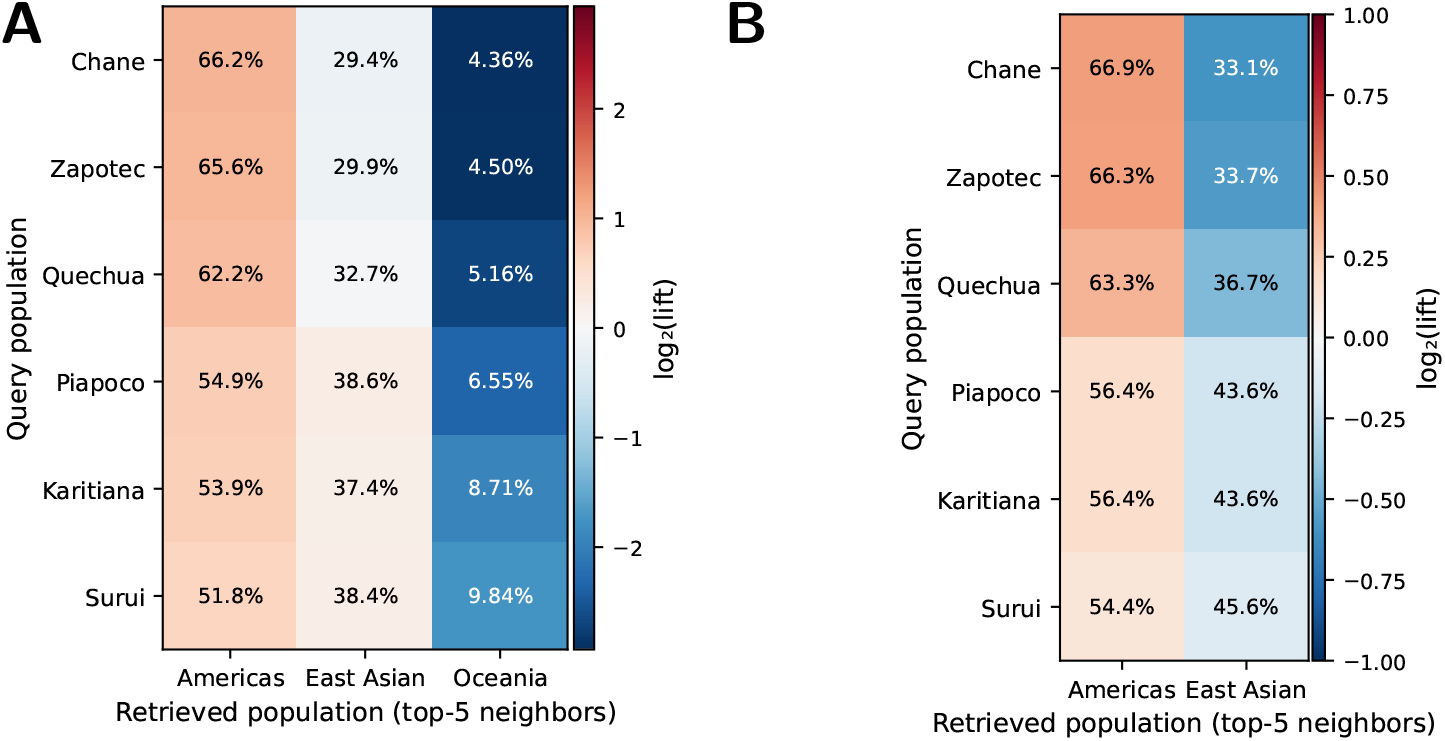
Population-scale affinities between the Americas, East Asia, and Oceania. **A**: For each query population we retrieve the top-5 nearest neighbors from the embedding corpus and tabulate the neighbors’ regional labels. Suruí and Karitiana show the highest enrichment for Oceanian retrieval. **B**: Negative-control retrieval with Oceanian references removed, recomputed over Americas and East Asian references only.

Across query populations, retrieved neighbors at the haplotype-level are predominantly labeled as Americas, with a consistent minority labeled as East Asian, matching the expected close genealogical relationship between the Americas and East Asia under models of the peopling of the Americas. In contrast, Oceania neighbors are rare overall, but show a marked relative increase in Suruí and Karitiana. Specifically, Suruí and Karitiana retrieve Oceania-labeled neighbors at 9.84% and 8.71%, respectively. The next highest is another Amazonian group, Piapoco, at 6.55%, compared with 4.36% to 5.16% in the other Indigenous American query populations (Chane, Zapotec, Quechua). This shift is reflected in the Oceania column’s log-lift shading, where Suruí and Karitiana exhibit the weakest depletion (closest to zero) relative to the background composition of the embedding corpus, consistent with a modest but detectable excess of Oceanian-reference nearest neighbors in these populations. As a robustness check, removing the Oceanian references shifts the retrieved neighbor composition primarily toward East Asian for Suruí, Karitiana, and Piapoco, whereas the other query populations remain dominated by neighbors in the Americas (Figure 5B).

Because each tokenized path corresponds to a marginal genealogy at a genomic interval, this enrichment can be read locally: a subset of Suruí and Karitiana segments have nearest-neighbor genealogical proximity to Oceanian reference paths that is stronger than in the other South American populations shown.

## 4. Conclusion

ARGformer leverages shared ancestral nodes to capture the topological intersections of the ARG. This allows the model to capture genealogical relationships across trees and the changes in ancestry induced by recombination along the genome. This representation serves as a flexible backbone for multiple downstream analyses, including reference-based settings in which newly threaded samples are approximately placed in an existing latent space and queried against nearby reference haplotypes. Through contrastive fine-tuning, the model learns to pull together related haplotypes and separate unrelated ones. This allows ARGformer to provide low-dimensional visualization of population structure from the ARG and to support local ancestry inference by classifying haplotypes in admixed individuals.

Together, the retrieval results support the interpretation that, although the dominant genealogical signal in South American genomes reflects shared Indigenous American ancestry with expected East Asian affinity, a subset of local genealogies in Suruí, Karitiana, and Piapoco populations are shared with ancestry found in Oceania. Similarly, we recovered expected signals of Denisovan introgression in Oceania. Because each tokenized path corresponds to a marginal tree along the genome, this retrieval-based view highlights how ARGformer can surface localized, ancestry-informative segments directly from inferred genealogies, without explicit genotype features, by exploiting the structure of local coalescent histories encoded in the ARG.

With appropriate architectural and objective modifications, we expect that the same framework can be extended to a broad range of other challenges in population genetics and evolutionary inference. The learned representations could potentially be adapted to graph-level tasks that detect demographic bottlenecks or signatures of selection (Browning & Browning, 2015; Vaughn & Nielsen, 2024). Scaling ARG-based models to biobank cohorts will also likely require more compressed representations. One promising direction is to define tokens that correspond to shared lineages in the genealogy, so that each individual is represented by a sequence of higher-level ancestral lineages over the past hundred or thousand generations, rather than by every individual ancestor. Future work could also explore decoder-only or encoder-decoder architectures for global graph-level tasks, while the present work focuses on encoder models and individual-level ancestry.

For ancestry inference, several limitations remain. Some paths in the marginal trees have few coalescent events, making them relatively uninformative and noisy for classification. A further limitation is ARG inference. Even the most scalable methods struggle with greater variant density that we expect in imputed data and whole-genomes (Gunnarsson et al., 2024). However, the breadth of population genetic information encoded in ARGs suggests that representation learning is a promising direction for future methods and applied analyses in population genetics.

## Acknowledgments

We thank Leo Speidel for helpful discussions and comments on this work.

## Supplementary Material

### S1 Dataset statistics

The real inferred ARG from (Wohns et al., 2022) includes samples from the 1000 Genomes Project (1KGP) (Auton et al., 2015), Simons Genome Diversity Project (SGDP) (Mallick et al., 2016), Human Genome Diversity Project (HGDP) (Bergström et al., 2020), three Neanderthal genomes (Chagyrskaya, Altai, and Vindija) (Mafessoni et al., 2020; Prüfer et al., 2014; 2017), and a Denisovan genome. We created two separate datasets for distinct analyses:

#### Archaic Introgression Analysis

This dataset includes archaic reference genomes (Altai, Chagyrskaya, and Vindija Neanderthals, and Denisovan), an African baseline (Yoruba), and held-out populations for validation (Utah Residents (CEPH) with Northern and Western European ancestry, Chinese Dai, Papuan Highlands, and Mandinka).

#### Oceania-like Ancestry in the Americas

This dataset consists of three reference groups and one test group: (1) East Asian populations (Han Chinese, Japanese), (2) Indigenous American baseline populations (Pima, Maya, Mixe), (3) Oceanian populations (Aboriginal Australian, Bougainville, Papuan populations), and a held-out test group of Indigenous American populations (Karitiana, Suruí, Chane, Piapoco, Quechua, Zapotec). Reference groups were balanced to the minimum group size to prevent biasing the model toward overrepresented populations.

We performed a 90%/10% train/validation split on the subset of unique marginal paths shown in Table S1.

**Table S1:** Dataset statistics for tasks on the real inferred ARG from (Wohns et al., 2022).

| Dataset | Statistic | Training | Validation |
| --- | --- | --- | --- |
| Archaic Introgression | Sample nodes | 688 |  |
|  | Vocabulary size | 184,963 |  |
|  | Unique paths | 7,660,071 | 851,119 |
|  | Total tokens | 120,297,364 | 13,369,481 |
| Oceania-like in Americas | Sample nodes | 665 |  |
|  | Vocabulary size | 119,067 |  |
|  | Unique paths | 3,473,898 | 385,989 |
|  | Total tokens | 57,777,684 | 6,421,781 |

### S2 Model training and efficiency

We adopted the ModernBERT architecture and optimization settings (Warner et al., 2025), using a 12-layer, 12-head transformer (Vaswani et al., 2017) with rotary position embeddings (RoPE) (Su et al., 2024), and Flash Attention 2 (Dao, 2024). We assessed the model’s ability to learn genealogical representations through self-supervised training. To measure generalization to unseen haplotypes, we monitored validation loss and masked token accuracy on a held-out set comprising 10% of ancestral paths. On the held-out set from the inferred ARG dataset, ARGformer achieved a top-1 accuracy of 95.96%, effectively reconstructing ancestors from context for previously unseen sequences. Training was also highly efficient. Pretraining sustained an average throughput of 189,500 tokens/sec and converged in 58 minutes on 1 NVIDIA A100 GPU with 80 GB of VRAM (Figure S1).

**Figure S1:**
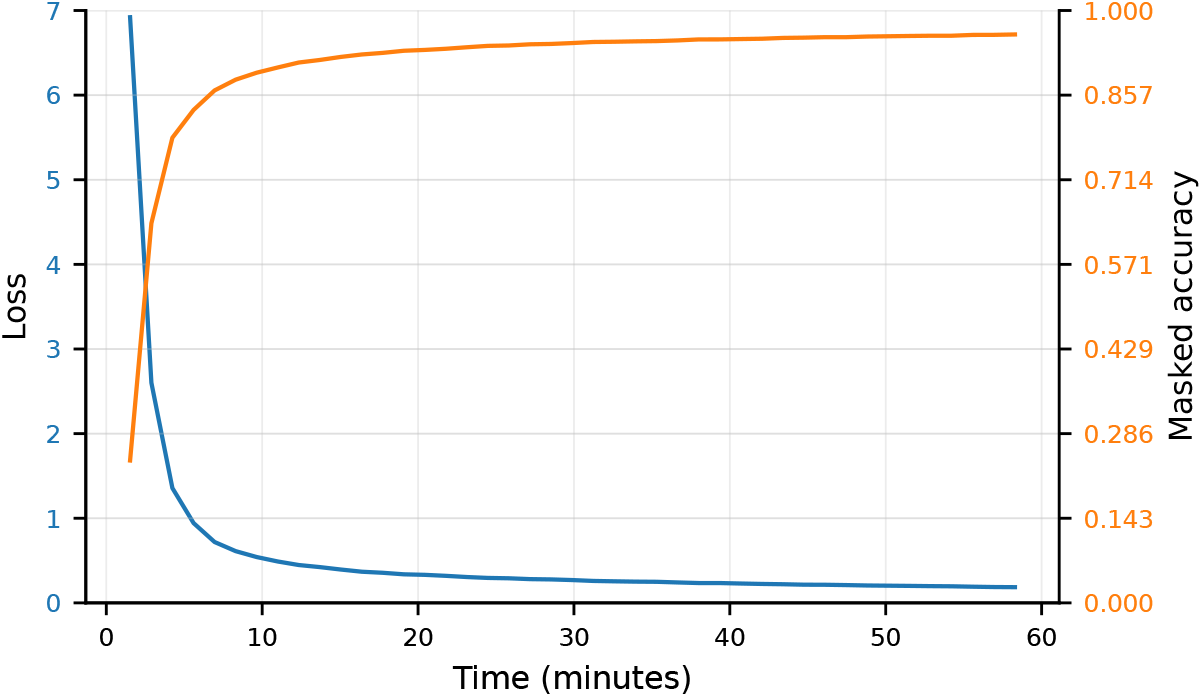
Training dynamics during self-supervised pretraining of ARGformer on the inferred ARG dataset. Validation loss decreases steadily while masked-token accuracy increases over time, showing stable convergence within a single 58 minute training run on one NVIDIA A100 GPU.

### S3 Visualization of Oceanian-like ancestry in South America

**Figure S2:**
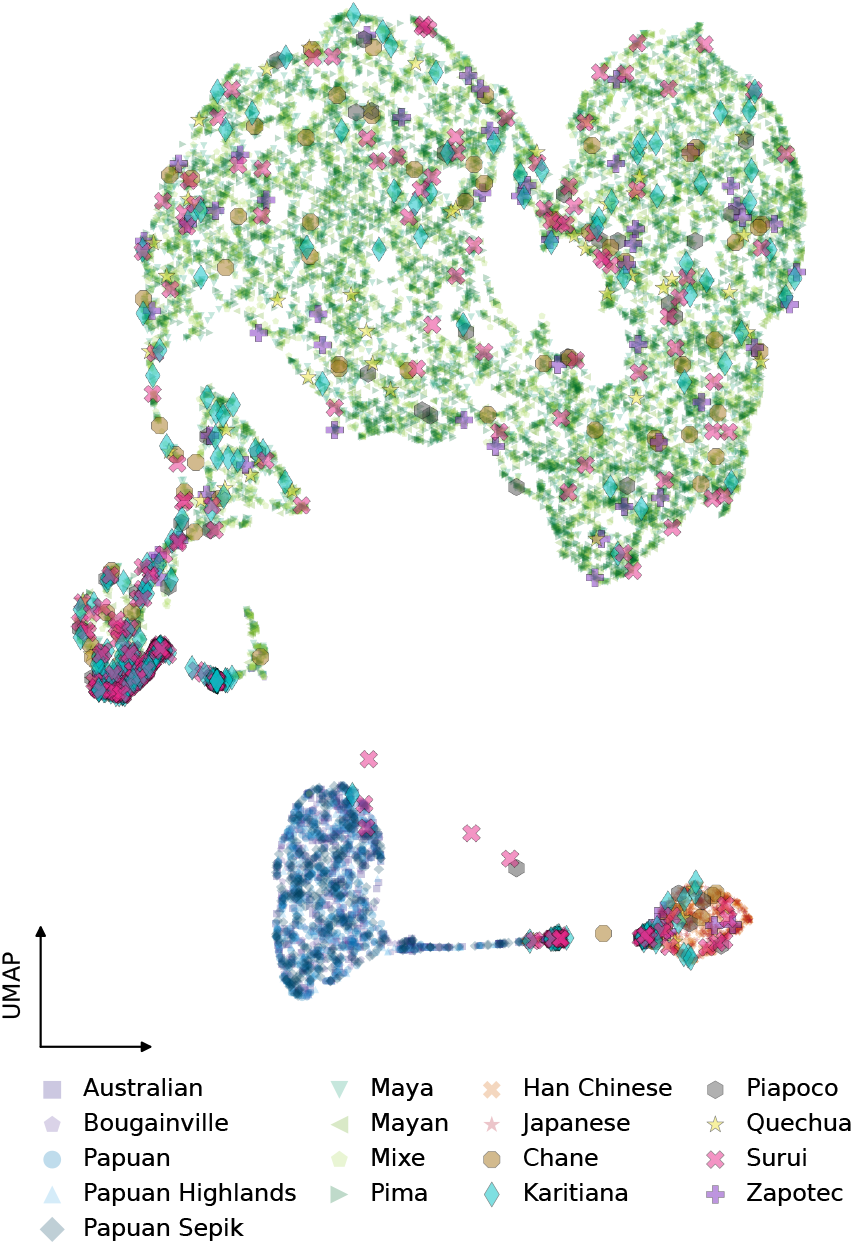
UMAP projection of haplotype-path embeddings for populations spanning Oceania (e.g., Australian, Bougainville, Papuan groups), East Asia (Han Chinese, Japanese), and the Americas (e.g., Chane, Karitiana, Piapoco, Quechua, Suruí, Zapotec).

A complementary view comes from a UMAP projection of the haplotype-path embedding space (Figure S2). Reference population groups form their own separated clusters with most of the query points spread across the American embedding manifold, meanwhile the Oceanian cluster contains a small set of Suruí and Karitiana embeddings. A small set of Suruí and Piapoco points are also displaced toward the Oceania side of the embedding space, consistent with the retrieval asymmetry observed above.

## S4 Analysis of learned representations

### 4.1. Additional probing analyses of genealogical structure in self-supervised embeddings

In Figure 3 we showed that a simple ridge probe trained on frozen self-supervised embeddings predicts the number of coalescent events along each path in both the American simulation and the archaic-modern inferred ARG, while collapsing under a permutation control. Here we provide an additional probe analysis for the Oceania-America inferred ARG, together with a brief summary of the probe setup.

For each dataset, we extracted frozen embeddings from the self-supervised model and trained a ridge regressor to predict the number of coalescent events along the corresponding leaf-to-root path. Evaluation was performed on held-out paths not used to fit the probe. As a null control, we repeated the same procedure after randomly permuting the training labels, which preserves the embedding distribution, probe class, and label marginal while breaking the correspondence between embeddings and their genealogical targets.

Figure S3 shows the same analysis in the Oceania-America inferred ARG. As in the main-text settings, predictions from the probe track the observed coalescent-event counts substantially better than the permutation control. Together with the American simulation and archaic-modern results shown in the main text, this supports the conclusion that self-supervised ARGformer embeddings encode genealogically meaningful information across simulated and real inferred genealogies, and across both continental-structure and archaic-introgression settings.

**Figure S3:**
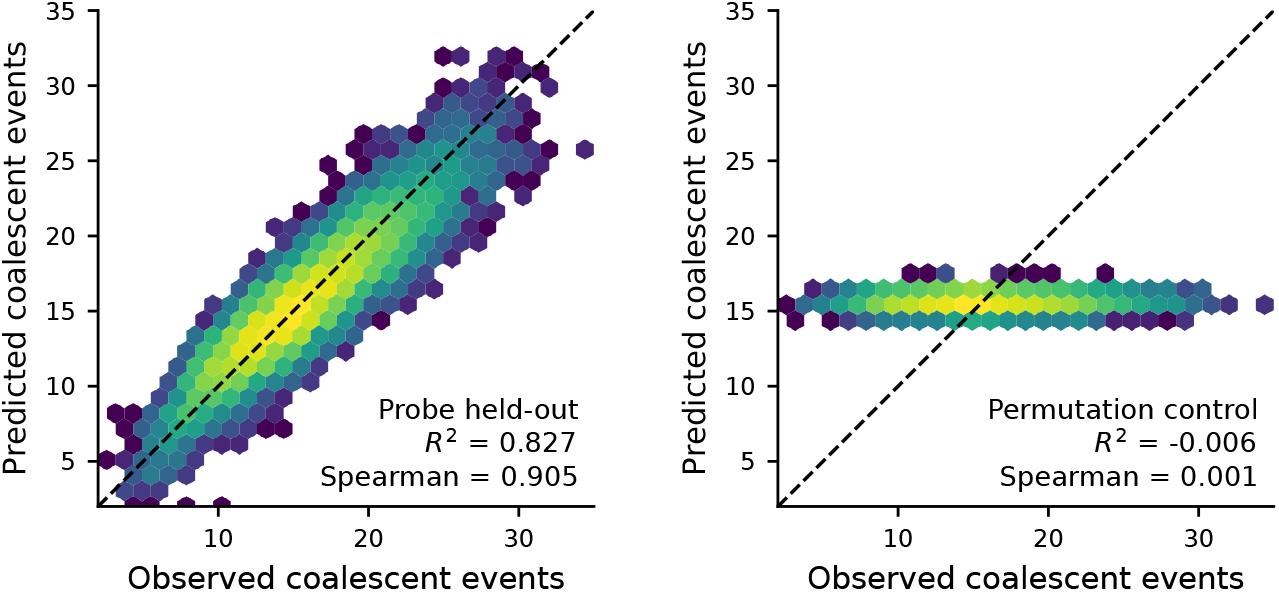
Additional probe analysis for the Oceania-America inferred ARG. The left plot shows held-out prediction of the number of coalescent events along each path from frozen self-supervised embeddings using a ridge probe, and the right plot shows the corresponding permutation control obtained by shuffling the training labels before fitting. As in the maintext analyses, the probe substantially outperforms the permutation control, indicating that the self-supervised embedding space encodes genealogical depth information beyond chance.

### 4.2. Attention patterns reflect structured genealogical relations

To examine what self-supervised pretraining learns beyond global embedding geometry, we analyzed node-to-node attention patterns on held-out paths from the American simulation. Figure S4 shows that different heads preferentially capture distinct genealogical relations along the leaf-to-root path. Representative examples include heads that attend to the immediate ancestor, heads that allocate attention toward the root, and heads that focus on deeper ancestors farther up the path.

These patterns are not uniformly distributed across the network. For each layer, we quantified each motif as the maximum fraction of node-to-node attention assigned to that pattern by any single head. The three behaviors peak in different layers, indicating that self-supervised pretraining does not simply produce a generic local-attention mechanism but instead yields heads with partially specialized genealogical roles. Immediate-ancestor attention is strongest early and in the last layers, while deep-ancestor attention is most sustained in the middle layers and drops sharply at the end. That fits with a mid-layer role of integrating higher-level genealogical context before the final layers refocus on local prediction.

**Figure S4:**
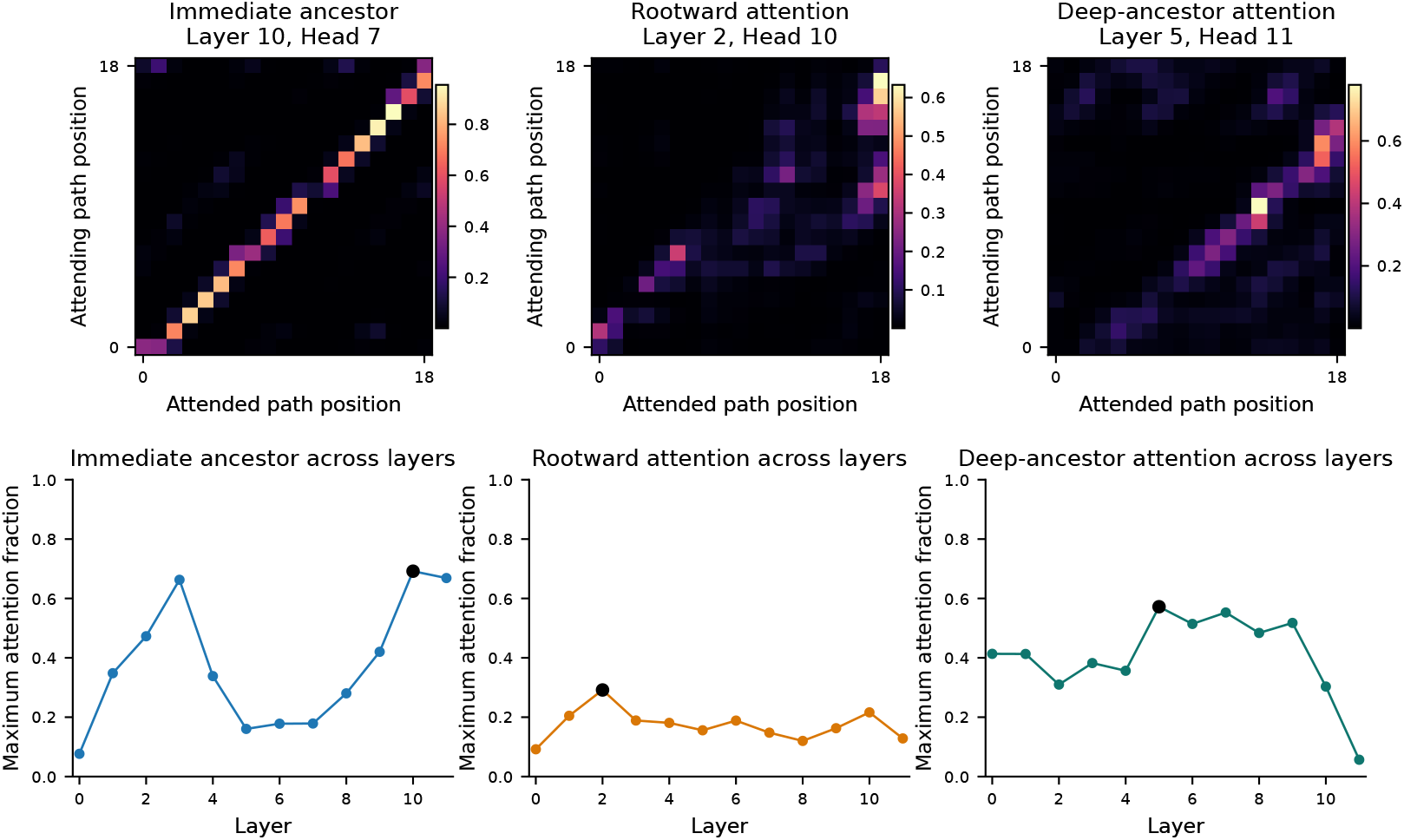
Attention interpretability analysis for held-out American simulation samples. Top row: representative attention maps for heads that preferentially attend to immediate ancestor (layer 10, head 7), rootward attention (layer 2, head 10), and deep-ancestor attention (layer 5, head 11). Bottom row: layer-wise progression of each pattern, quantified as the maximum fraction of node-to-node attention assigned to that pattern by any single head in each layer. These panels show that different heads preferentially capture distinct genealogical patterns rather than all collapsing onto the same local signal.

## References

Jeffrey R. Adrion, Christopher B. Cole, Noah Dukler et al. A community-maintained standard library of population genetic models. eLife, 9:e54967, 2020.

Hussein Al-Asadi, Desislava Petkova, Matthew Stephens et al. Estimating recent migration and population-size surfaces. PLOS Genetics, 15(1):e1007908, 2019.

Adam Auton, Gonçalo R. Abecasis, David M. Altshuler et al. A global reference for human genetic variation. Nature, 526(7571):68–74, 2015.

C. J. Battey, Gabrielle C. Coffing, and Andrew D. Kern. Visualizing population structure with variational autoencoders. G3, 11(1):jkaa036, 2021.

Anders Bergström, Shane A. McCarthy, Ruoyun Hui et al. Insights into human genetic variation and population history from 929 diverse genomes. Science, 367(6484):eaay5012, 2020.

Sharon R. Browning and Brian L. Browning. Accurate non-parametric estimation of recent effective population size from segments of identity by descent. The American Journal of Human Genetics, 97(3):404–418, 2015.

Sharon R. Browning, Brian L. Browning, Martha L. Daviglus et al. Ancestry-specific recent effective population size in the Americas. PLOS Genetics, 14(5):e1007385, 2018.

Sharon R. Browning, Ryan K. Waples, and Brian L. Browning. Fast, accurate local ancestry inference with FLARE. American Journal of Human Genetics, 110(2):326–335, 2023.

Clare Bycroft, Colin Freeman, Desislava Petkova et al. The UK Biobank resource with deep phenotyping and genomic data. Nature, 562(7726):203–209, 2018.

Marcos Araújo Castro e Silva, Tiago Ferraz, Maria Cátira Bortolini et al. Deep genetic affinity between coastal pacific and amazonian natives evidenced by australasian ancestry. Proceedings of the National Academy of Sciences of the USA, 118(14):e2025739118, 2021.

Tri Dao. Flashattention-2: Faster attention with better parallelism and work partitioning. In ICLR, 2024.

Yun Deng, Rasmus Nielsen, and Yun S. Song. Robust and accurate Bayesian inference of genome-wide genealogies for hundreds of genomes. Nature Genetics, pp. 1–12, 2025.

Jacob Devlin, Ming-Wei Chang, Kenton Lee et al. BERT: Pre-training of deep bidirectional transformers for language understanding. In Proceedings of the 2019 Conference of the North American Chapter of the Association for Computational Linguistics: Human Language Technologies, pp. 4171–4186, 2019.

Alex Diaz-Papkovich, Luke Anderson-Trocmé, Chief Ben-Eghan et al. UMAP reveals cryptic population structure and phenotype heterogeneity in large genomic cohorts. PLOS Genetics, 15(11):e1008432, 2019.

Lex Flagel, Yaniv Brandvain, and Daniel R. Schrider. The unreasonable effectiveness of convolutional neural networks in population genetic inference. Molecular Biology and Evolution, 36(2):220–238, 2019.

Margarita Geleta, Daniel Mas Montserrat, Xavier Giro-i Nieto et al. Autoencoders for genomic variation analysis. Genome Research, 36(2):348–360, 2026.

Robert C. Griffiths and Paul Marjoram. An ancestral recombination graph. Progress in Population Genetics and Human Evolution, pp. 257–270, 1997.

Árni Freyr Gunnarsson, Jiazheng Zhu, Brian C. Zhang et al. A scalable approach for genome-wide inference of ancestral recombination graphs. bioRxiv, 2024.

Zhenyu Hou, Xiao Liu, Yukuo Cen et al. GraphMAE: Self-supervised masked graph autoencoders. In Proceedings of the 28th ACM SIGKDD Conference on Knowledge Discovery and Data Mining, pp. 594–604, 2022.

Richard R. Hudson. Properties of a neutral allele model with intragenic recombination. Theoretical Population Biology, 23(2):183–201, 1983.

Guy S. Jacobs, Georgi Hudjashov, Lauri Saag et al. Multiple deeply divergent Denisovan ancestries in Papuans. Cell, 177(4):1010–1021.e32, 2019.

Yanrong Ji, Zhihan Zhou, Han Liu et al. DNABERT: pre-trained bidirectional encoder representations from transformers model for DNA-language in genome. Bioinformatics, 37 (15):2112–2120, 2021.

Jerome Kelleher, Alison M. Etheridge, and Gilean McVean. Efficient coalescent simulation and genealogical analysis for large sample sizes. PLOS Computational Biology, 12(5):e1004842, 2016.

Jerome Kelleher, Yan Wong, Anthony W. Wohns et al. Inferring whole-genome histories in large population datasets. Nature Genetics, 51(9):1330–1338, 2019.

Jinwoo Kim, Dat Nguyen, Seonwoo Min et al. Pure transformers are powerful graph learners. Advances in Neural Information Processing Systems, 35:14582–14595, 2022.

Kevin Korfmann, Nathaniel S Pope, Melinda Meleghy et al. Coalescence and translation: A language model for population genetics. bioRxiv, 2025.

M. Elise Lauterbur, Maria Izabel A. Cavassim, Ariella L. Gladstein et al. Expanding the Stdpopsim species catalog, and lessons learned for realistic genome simulations. eLife, 12, 2023.

Alexander L. Lewanski, Michael C. Grundler, and Gideon S. Bradburd. The era of the ARG: An introduction to ancestral recombination graphs and their significance in empirical evolutionary genomics. PLOS Genetics, 20(1):e1011110, 2024.

Fabrizio Mafessoni, Steffi Grote, Cesare de Filippo et al. A high-coverage Neandertal genome from Chagyrskaya Cave. Proceedings of the National Academy of Sciences, 117 (26):15132–15136, 2020.

Swapan Mallick, Heng Li, Mark Lipson et al. The Simons Genome Diversity Project: 300 genomes from 142 diverse populations. Nature, 538(7624):201–206, 2016.

Gil McVean. A genealogical interpretation of principal components analysis. PLOS Genetics, 5(10):e1000686, 2009.

Rasmus Nielsen, Andrew H. Vaughn, and Yun Deng. Inference and applications of ancestral recombination graphs. Nature Reviews Genetics, 26(1):47–58, 2025.

Aaron van den Oord, Yazhe Li, and Oriol Vinyals. Representation learning with contrastive predictive coding. arXiv preprint 1807.03748, 2018.

Nick Patterson, Alkes L. Price, and David Reich. Population structure and eigenanalysis. PLOS Genetics, 2(12):e190, 2006.

Kay Prüfer, Fernando Racimo, Nick Patterson et al. The complete genome sequence of a Neanderthal from the Altai Mountains. Nature, 505(7481):43–49, 2014.

Kay Prüfer, Cesare de Filippo, Steffi Grote et al. A high-coverage Neandertal genome from Vindija Cave in Croatia. Science, 358(6363):655–658, 2017.

Matthew D. Rasmussen, Melissa J. Hubisz, Ilan Gronau et al. Genome-wide inference of ancestral recombination graphs. PLOS Genetics, 10(5):e1004342, 2014.

Alexander Rives, Stephan Meier, Tom Sercu et al. Biological structure and function emerge from scaling unsupervised learning to 250 million protein sequences. Proceedings of the National Academy of Sciences of the USA, 118(15):e2016239118, 2021.

Daniel R. Schrider and Andrew D. Kern. Supervised machine learning for population genetics: A new paradigm. Trends in Genetics, 34(4):301–312, 2018.

Sara Sheehan and Yun S. Song. Deep learning for population genetic inference. PLOS Computational Biology, 12(3):e1004845, 2016.

Pontus Skoglund, Swapan Mallick, Maria Cátira Bortolini et al. Genetic evidence for two founding populations of the Americas. Nature, 525(7567):104–108, 2015.

Leo Speidel, Marie Forest, Sinan Shi et al. A method for genome-wide genealogy estimation for thousands of samples. Nature Genetics, 51(9):1321–1329, 2019.

Jianlin Su, Murtadha Ahmed, Yu Lu et al. Roformer: Enhanced transformer with rotary position embedding. Neurocomputing, 568:127063, 2024.

Ashish Vaswani, Noam Shazeer, Niki Parmar et al. Attention is all you need. NeurIPS, 30, 2017.

Andrew H. Vaughn and Rasmus Nielsen. Fast and accurate estimation of selection coefficients and allele histories from ancient and modern DNA. Molecular Biology and Evolution, 41 (8):msae156, 2024.

Davide M. Vespasiani, Guy S. Jacobs, Laura E. Cook et al. Denisovan introgression has shaped the immune system of present-day papuans. PLOS Genetics, 18(12):e1010470, 2022.

Benjamin Warner, Antoine Chaffin, Benjamin Clavié et al. Smarter, better, faster, longer: A modern bidirectional encoder for fast, memory efficient, and long context finetuning and inference. In Proceedings of the 63rd Annual Meeting of the Association for Computational Linguistics, pp. 2526–2547, 2025.

Anthony Wilder Wohns, Yan Wong, Ben Jeffery et al. A unified genealogy of modern and ancient genomes. Science, 375(6583):eabi8264, 2022.

Yan Wong, Anastasia Ignatieva, Jere Koskela et al. A general and efficient representation of ancestral recombination graphs. Genetics, 228(1):iyae100, 2024.

Brian C. Zhang, Arjun Biddanda, Árni Freyr Gunnarsson et al. Biobank-scale inference of ancestral recombination graphs enables genealogical analysis of complex traits. Nature Genetics, 55(5):768–776, 2023.

